# Hippocampal neural fluctuation between memory encoding and retrieval states during a working memory task in humans

**DOI:** 10.1101/2023.04.28.538785

**Authors:** Yusuke Watanabe, Yuji Ikegaya, Takufumi Yanagisawa

## Abstract

**Background:** Working memory (WM) is essential for everyday life, yet its neural mechanism remains unclear. Although the hippocampus plays a critical role in memory consolidation and retrieval, its role in WM tasks has yet to be fully elucidated. We hypothesized that multiunit activities in the hippocampus alter their representations depending on the memory load and phase of a WM task, particularly in relation to sharp-wave ripple complexes (SWRs), which are known as various cognitive biomarkers.

**Methods:** We used an open dataset of intracranial electroencephalogram (iEEG) data and multiunit activity recorded from the medial temporal lobe (MTL) of nine patients with epilepsy. The MTL includes the hippocampus, entorhinal cortex, and amygdala. During the recording, all subjects performed an eight-second Sternberg test, in which they memorized sets of four, six, or eight letters (encoding phase), waited for three seconds (maintenance phase), and recalled whether a probe letter was included (Match IN task) or not (Mismatch OUT task) (retrieval phase). We used Gaussian-process factor analysis to visualize the neural trajectories of multiunit activity in MTL regions during the task. We also detected SWRs from the iEEG data in the hippocampus.

**Findings:** We found that the trajectory distance between phases of the Sternberg task was larger in the hippocampus compared to the entorhinal cortex and amygdala. Additionally, the trajectory distance between the encoding and retrieval phases was memory load dependent. Moreover, a transient trajectory increase was detected during SWRs. Finally, the trajectory direction of the hippocampus fluctuated between the encoding and retrieval states, and the balance of the fluctuation was shifted to the retrieval state during SWR periods.

**Interpretation:** Our results demonstrate the involvement of the hippocampus during a WM task. Furthermore, it is suggested that SWR in the retrieval phase plays a role in memory retrieval for a WM task. Our results provide new insight into the two-stage model of memory formation.

## 1 Introduction

Working memory (WM) is crucial in everyday life; however, its neural mechanism has yet to be fully elucidated, and the role of the hippocampus in WM tasks remains unclear. For example, a famous patient – H.M. – who had his hippocampus removed did not show WM impairments (Scoville & Milner, 1957) and retained numbers for over 15 minutes by continuous rehearsal (Squire, 2009). Recent studies, however, have shown that the hippocampus is involved in WM tasks, especially during the maintenance period, as indicated by persistent spiking (Boran et al., 2019; Jan Kamiński et al., 2017; Kornblith et al., 2017; Faraut et al., 2018) and increased gamma oscillation power (van Vugt et al., 2010). Moreover, WM performance depends on coordinating local field potential (LFP) power and delta oscillations in the hippocampus (Leszczyński et al., 2015).

Among the hippocampal phenomena, a transient and synchronous oscillation called sharp-wave ripple (SWR; Buzsáki, 2015) is associated with various cognitive functions, including memory replay (Wilson and McNaughton, 1994; Nádasdy et al., 1999; Lee and Wilson, 2002; Diba, K., & Buzsáki, G, 2007; Davidson et al., 2009), memory consolidation (Girardeau et al., 20009; Ego-Stengel et al., 2010; Fernández-Ruiz et al., 2019; Kim et al., 2022), recall (Wu et al., 2017; Norman et al., 2019; Norman et al., 2021), and neural plasticity (Behrens et al., 2005; Norimoto et al., 2018). Thus, SWR might be a fundamental observation reflecting the information processing of the hippocampus, although its role in WM remains unclear (Jadhav et al., 2012; Fernández-Ruiz et al., 2019).

Hippocampal neurons may exhibit low-dimensional representations during WM tasks. For instance, firing patterns of place cells (O’Keefe and Dostrovsky, 1971; O’Keefe, 1976; Ekstrom et al., 2003; Kjelstrup et al., 2008; Harvey et al., 2009) in the hippocampus were embedded into a dynamic, nonlinear 3D hyperbolic geometry while navigating (Zhang et al., 2022). Additionally, grid cells in the entorhinal cortex (EC), which is the main gate of the hippocampus (Naber et al., 2001; van Strien et al., 2009; Strange et al., 2014), showed toroidal topology during exploration (Gardner et al., 2022). In addition to navigation tasks, the hippocampus is expected to show low-dimensional latent spaces for WM tasks.

One of the challenges in studying hippocampal functions in WM tasks is the lack of appropriate experimental designs. In nonhuman animals, WM tasks are often trained through multiple trials using maze or classical conditioning methods, which do not distinguish between the acquisition and recall of information. In humans, noninvasive recording methods such as electroencephalography (EEG) and functional magnetic resonance imaging (fMRI) have limited spatial and temporal resolution. Invasive recordings of deep brain regions, such as intracranial EEG (iEEG), provide higher noise-to-ratio signals but limit the number of experiments and pose challenges in the histological verification of recording sites.

In this study, we investigated the hypothesis that hippocampal neurons express distinct representations as trajectories in low-dimensional spaces during a WM task with a specific focus on SWR periods. To test this hypothesis, we used a dataset of patients performing a WM task with high temporal resolution (1 s for fixation, 2 s for encoding, 3 s for maintenance, and 2 s for retrieval). Simultaneously, their iEEG data were recorded with tetrodes in the medial temporal lobe (MTL) regions. We used Gaussian-process factor analysis (GPFA; Yu et al., 2009) on the multiunit activity to calculate low-dimensional neural trajectories in the MTL.

## 2 Methods

### 2.1 Dataset

A publicly available dataset was employed, in which nine subjects performed a modified Sternberg task that consisted of fixation (1 s), encoding (2 s), maintenance (3 s), and retrieval (2 s) phases (Boran et al., 2020). Subjects were presented with four, six, or eight alphabetical letters in the encoding phase. They were expected to recall whether a probe letter in the retrieval phase was also displayed (Match IN) or was not (Mismatch OUT). iEEG signals were recorded at 32 kHz (0.5– 5,000 Hz passband) during the modified Sternberg task with depth electrodes in the medial temporal (MTL) regions (*i.e.*, left and right hippocampal head [AHL and AHR], hippocampal body [PHL and PHR], EC [ECL and ECR], and amygdala [AL and AR]; Figure 1A & Table 1). iEEG signals were resampled at 2 kHz. Correlations were found among the experimental variables (Fig. S1). The timings of multiunit spikes were estimated by a spike sorting algorithm (Niediek et al., 2016) by the Combinato package (https://github.com/jniediek/combinato) from the 32-kHz LFP signals (Figure 1C).

**Figure 1.**
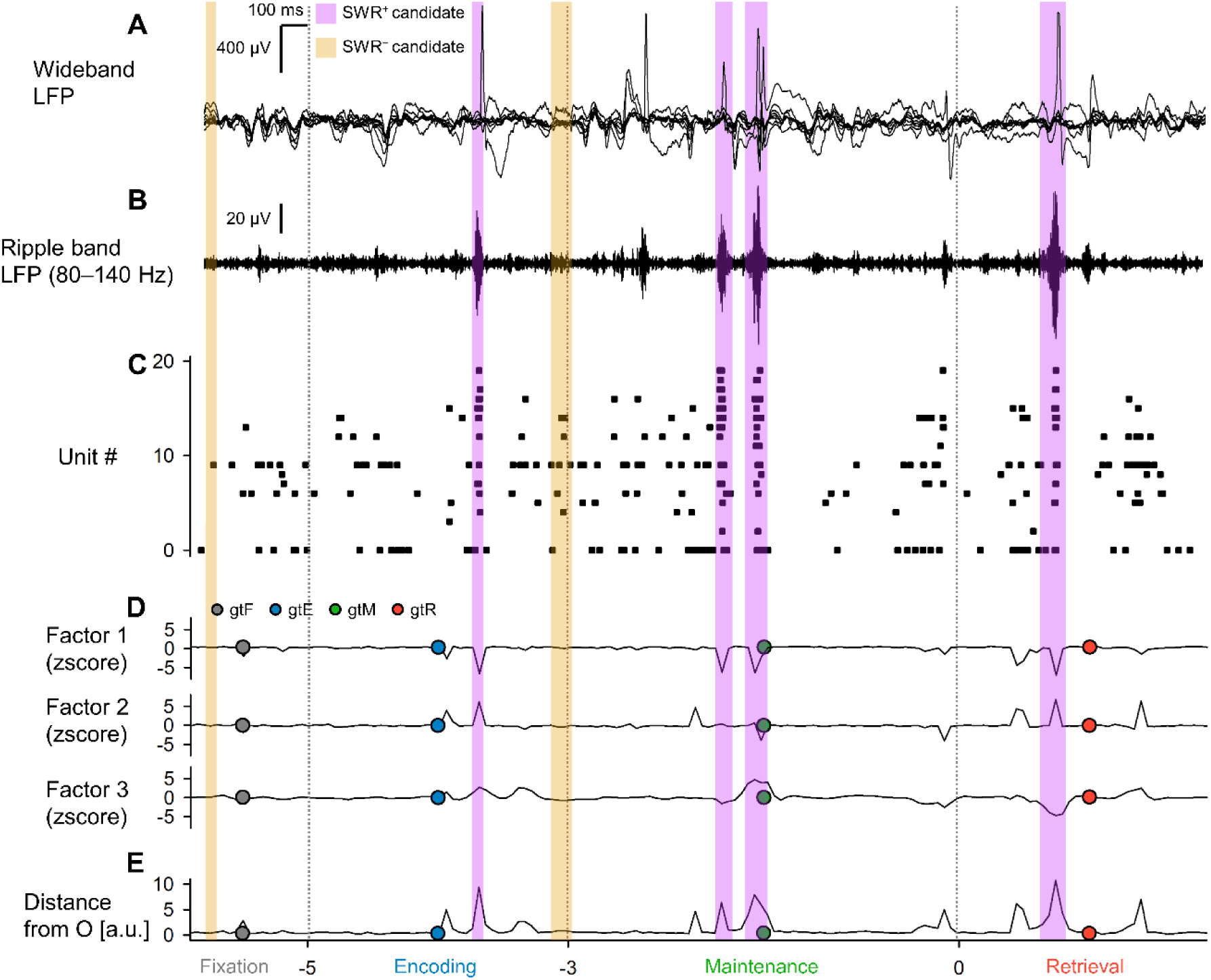
Representative LFP trace and neural trajectory of the hippocampus during a working memory task. ***A–B.*** Representative local field potential (LFP) traces of intracranial encephalogram (iEEG) data (subject #6, session #2, trial #5, in the left hippocampal head) in the wideband (***A***) and ripple bands (***B***) recorded with 8-channel tetrodes during a modified Sternberg working memory task (*gray*, fixation, 1 s; *blue,* encoding, 2 s; *green*, maintenance, 3 s; and *red*, retrieval phase, 2 s). ***C.*** Raster plot of multiunit spikes for the same trial. ***D***. The first three factors of hippocampal trajectory in the same trial calculated by Gaussian-process factor analysis on spike counts for 50- ms bins. Dot circles show the coordinates of trial geometric medians for the four phases. ***E***. Trajectory distance of the first three factors from the origin (O). Note that superimposed rectangles show the timings of SWR^+^ candidates (*purple*) and SWR^−^ candidates (*yellow*).

**Table 1.**
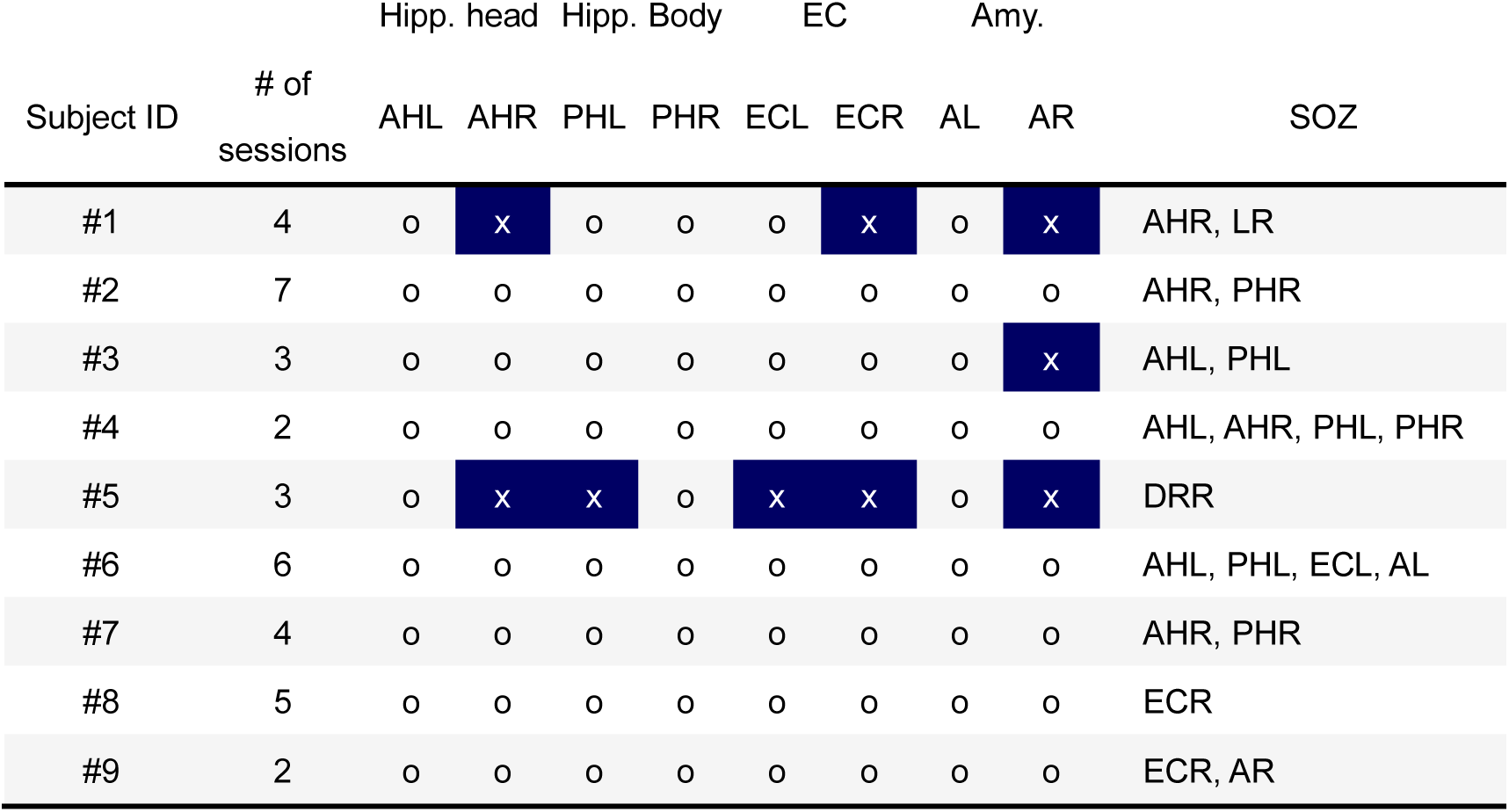
Electrode positions of the dataset. The electrode positions and the seizure onset zones. Regions marked as “o” were available, but those marked as “x” (*Navy*) were not available in the dataset. Abbreviations: AHL, left hippocampal head; AHR, right hippocampal head; PHL, left hippocampal body; PHR, right hippocampal body; ECL, left entorhinal cortex; ECR, right entorhinal cortex; AL, left amygdala; AR, right amygdala, SOZ: seizure onset zone.

| Subject ID | # of sessions | Hipp. head |  | Hipp. Body |  | EC |  | Amy. |  | SOZ |
| --- | --- | --- | --- | --- | --- | --- | --- | --- | --- | --- |
|  |  | AHL | AHR | PHL | PHR | ECL | ECR | AL | AR |  |
| #1 | 4 | o | x | o | o | o | x | o | x | AHR, LR |
| #2 | 7 | o | o | o | o | o | o | o | o | AHR, PHR |
| #3 | 3 | o | o | o | o | o | o | o | x | AHL, PHL |
| #4 | 2 | o | o | o | o | o | o | o | o | AHL, AHR, PHL, PHR |
| #5 | 3 | o | x | x | o | x | x | o | x | DRR |
| #6 | 6 | o | o | o | o | o | o | o | o | AHL, PHL, ECL, AL |
| #7 | 4 | o | o | o | o | o | o | o | o | AHR, PHR |
| #8 | 5 | o | o | o | o | o | o | o | o | ECR |
| #9 | 2 | o | o | o | o | o | o | o | o | ECR, AR |

### 2.2 Neural trajectory calculation by Gaussian-process factor analysis

GPFA (Yu et al., 2009) was applied to the multiunit activity by session to determine neural trajectories (factors; Figure 1D) of trials in the hippocampus, EC, and amygdala (Figure 1D). GPFA was implemented by the Python elephant package (https://elephant.readthedocs.io/en/latest/reference/gpfa.html). The bin size was set as 50 ms. Each factor was z-normalized by session. From trajectories, the Euclidean distance from the origin (O (0,0,0)) (Figure 1E) was calculated.

From every trajectory at a region (*e.g.*, AHL), “geometric medians” during the four phases were calculated (*i.e.*, gF, gE, gM, and gR during the fixation, encoding, maintenance, and retrieval phases, respectively) by taking the medians of trajectory coordinates during the phase (Figure 1D). The optimal dimension for GPFA was determined as three; this was calculated via the elbow method on the log-likelihood values and evaluated by the threefold cross-validation method (Figure 2B).

**Figure 2.**
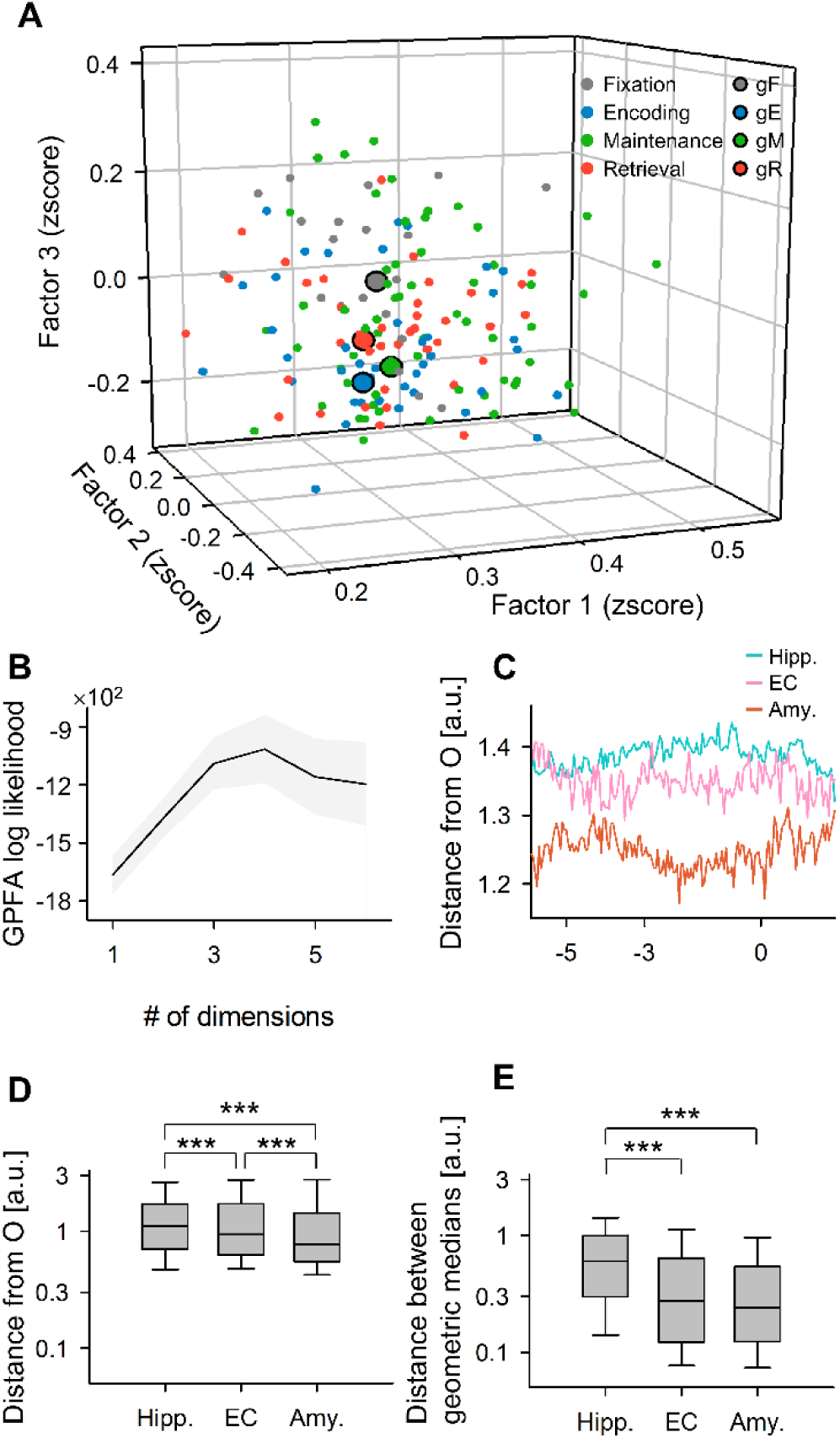
State-dependent hippocampal neural trajectory. ***A.*** A representative neural trajectory of the left hippocampus calculated by Gaussian-process factor analysis (GPFA). Small points (*n* = 160) show coordinates of 50-ms bins of neural trajectory in a session (median of 50 trials). *Black*-edged large points show the trajectory geometric medians of the four phases of a Sternberg working memory task (*gray*, fixation, 1 s; *blue,* encoding, 2 s; *green*, maintenance, 3 s; and *red*, retrieval phase, 2 s). ***B.*** Log-likelihoods of predictions of GPFA models as a function of the number of dimensions to embed (mean ± 95% confidence interval). Note that the optimal dimension was determined to be three via the elbow method. ***C***. Trajectory distance from the origin (O) in the hippocampus (Hipp.), entorhinal cortex (EC), and amygdala (Amy.), as a function of time from probe (mean ± 95% confidential interval). ***D.*** The trajectory distances from O in medial temporal regions were greater in the following order: Hipp. (1.11 [1.01]; median [IQR]; *n* = 195.681 time points), EC (0.94 [1.10]; median [IQR]; *n* = 133,761 time points), and Amy. (0.78 [0.88]; median [IQR]; *n* = 165,281 time points). ***E***. Distances among trial trajectory geometric medians of phases (*i.e.*, d(gtF, gtE), d(gtF, gtM), d(gtF, gtR), d(gtE, gtM), d(gtE, gtR), d(gtM, gtR)) were pooled and longer in the Hipp. (0.60 [0.70]; median [IQR]; *n* = 8,772 combinations) than in EC (0.28 [0.52]; median [IQR]; *n* = 5,017 combinations; *p* < 0.01; Brunner–Munzel test) or Amy. (0.24 [0.42]; median [IQR]; *n* = 7,466 combinations; *p* < 0.01; Brunner–Munzel test).

### 2.3 Defining hippocampal SWR candidates from hippocampal regions

Hippocampal SWR candidates were detected based on a consensus constructed by researchers (Liu et al., 2022) as follows. LFP signals of a region of interest (ROI; *e.g.*, AHL) were rereferenced by subtracting a control signal calculated by averaging out-of-ROI signals (*e.g.*, AHR, PHL, PHR, ECL, ECR, AL, and AR; Figure 1A). SWR candidates (= SWR^+^ candidates) were defined from the LFP signals passed through the ripple band filter (80–140 Hz) (Figure 1B) using a published tool (https://github.com/Eden-Kramer-Lab/ripple_detection; Kay et al., 2016) with modifications such as updated ripple band range from 150–250 (for rodents) to 80–140 Hz (for humans).

As control events of SWR^+^ candidates, SWR^−^ candidates were defined by shuffling the timestamps of SWR^+^ candidates across all trials from all subjects. SWR^+^/SWR^−^ candidates were visually inspected (Figure 1).

### 2.4 Defining hippocampal SWRs from putative CA1 regions

Putative CA1 regions were defined as follows. SWR^+^/SWR^−^ candidates were embedded into a two-dimensional space based on their superimposed spike counts per unit using UMAP (uniform manifold approximation and projection; McInnes et al., 2018) in a supervised fashion (Figure 4A). The silhouette score (Rousseeuw et al., 1987), a validation barometer for clustering, was calculated from clustered samples (Table 2). The hippocampal regions with silhouette scores greater than 0.6 on average across sessions (75^th^ percentile) (Figure 4B) were defined as putative CA1 regions (Table 3; *i.e.*, AHL of subject #1, AHR of subject #3, PHL of subject #4, AHL of subject #6, and AHR of subject #9). SWR^+^/SWR^−^ candidates in putative CA1 regions were defined as SWR^+^/SWR^−^ (no longer candidates). The duration [ms] and ripple band peak amplitude [SD of baseline] of detected SWRs were calculated (Figure 4C & E). SWR^+^/SWR^−^ were visually inspected as shown in Figure 1.

**Figure 4.**
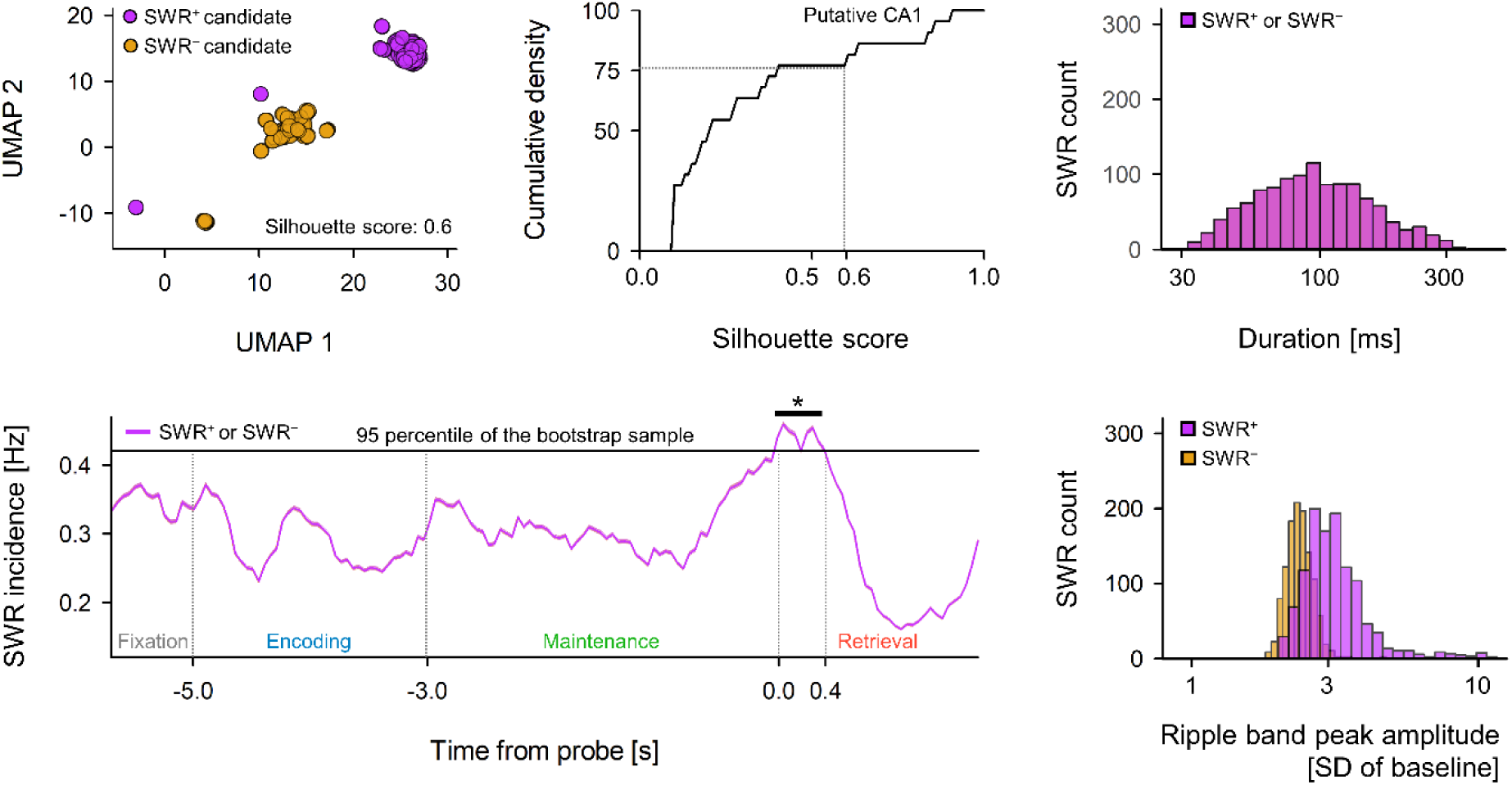
SWR detection from putative CA1 regions. ***A***. A representative embedding of unit activities during SWR^+^ candidates (*purple*; candidates of SWR event) and SWR^−^ candidates (*yellow*; control of SWR^+^ candidates) using two-dimensional UMAP (uniform manifold approximation and projection; Mclnnes et al., 2018). ***B***. Cumulative density plot of the silhouette scores of hippocampal regions (Table 2) as a clustering barometer. Note that hippocampal regions with silhouette scores greater than 0.60 (75^th^ percentile) were defined as putative CA1 regions, and from putative CA1 regions, SWR^+^ candidates and SWR^−^ candidates were defined as SWR^+^ (*n* = 1,170) and SWR^−^ (*n* = 1,170), respectively. ***C.*** Distribution of duration [ms] of SWR^+^ (*purple*; 93.0 [65.4], median [IQR]) and SWR^−^ (*yellow*; 93.0 [65.4], median [IQR]). Note that the two distributions exactly overlapped because of their definitions. ***D***. Ripple incidence [Hz] of SWR^+^ (*purple*) and SWR^−^ (*yellow*) as a function of time from probe (mean ± 95% confidential interval). Note that SWR incidence at 0–400 ms from the probe was significantly larger than the 95^th^ percentile of the bootstrap samples (0.421 [Hz], \**p* < 0.05). ***E.*** Distribution of ripple band peak amplitude [SD of baseline] of SWR^−^ (*yellow*; 2.37 [0.33], median [IQR]) and SWR^+^ (*purple*; 3.05 [0.85], median [IQR]) (\*\*\**p* < 0.001; Brunner–Munzel test).

**Table 2.** The silhouette score of UMAP clustering between SWR^+^ candidates and SWR^−^candidates. The silhouette scores (mean ± SD for sessions by subject) of UMAP clustering on SWR^+^ candidates and SWR^−^ candidates (Figure 4A) were based on their underlying multiunit spike patterns (mean values were 0.205 [0.285], median [IQR]; Figure 4B).

| Subject | Hipp. head |  | Hipp. body |  |
| --- | --- | --- | --- | --- |
|  | AHL | AHR | PHL | PHR |
| #1 | 0.60 ± 0.14 | n.a. | n.a. | 0.1 ± 0 |
| #2 | 0.21 ± 0.16 | 0.17 ± 0.21 | 0.18 ± 0.22 | 0.20 ± 0.15 |
| #3 | 0.40 ± 0.42 | 0.83 ± 0.12 | n.a. | n.a. |
| #4 | 0.10 ± 0.00 | 0.10 ± 0.00 | 0.90 ± 0.00 | 0.10 ± 0.14 |
| #5 | n.a. | n.a. | n.a. | n.a. |
| #6 | 0.63 ± 0.06 | n.a. | n.a. | 0.27 ± 0.06 |
| #7 | 0.10 ± 0.00 | 0.35 ± 0.35 | 0.37 ± 0.47 | 0.10 ± 0.00 |
| #8 | 0.13 ± 0.10 | n.a. | 0.28 ± 0.49 | n.a. |
| #9 | n.a. | 0.85 ± 0.07 | 0.15 ± 0.07 | n.a. |

**Table 3.**
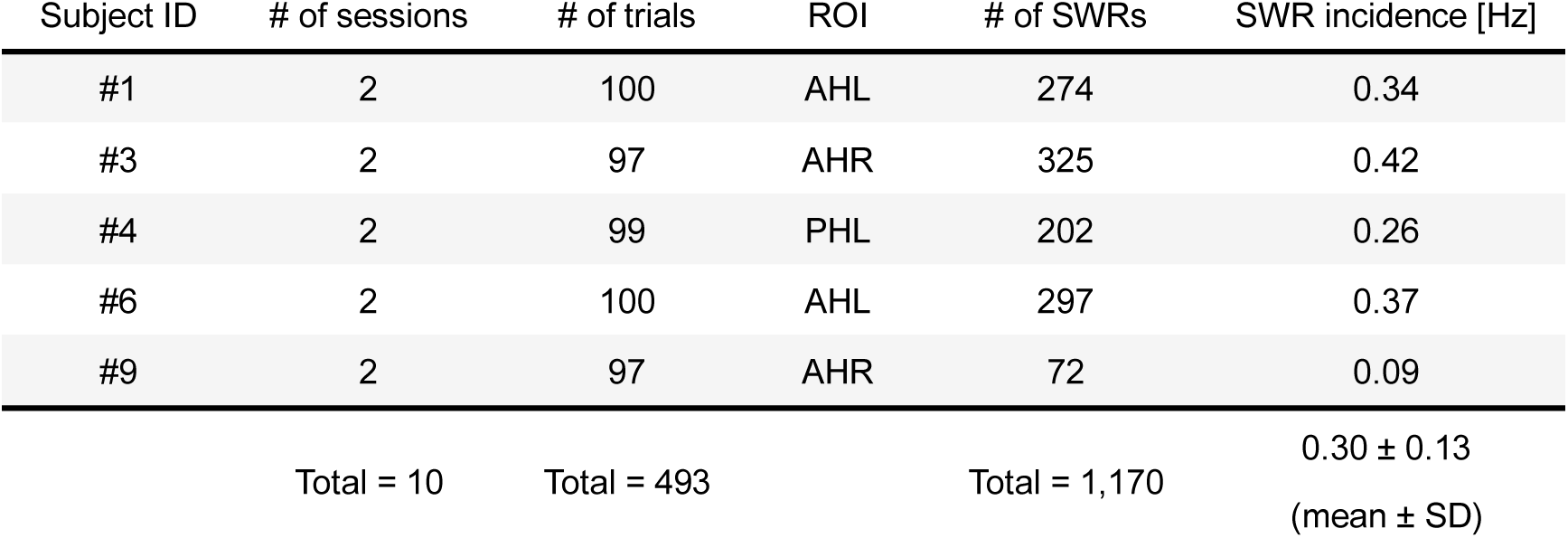
The number of defined SWR events. The table summarizes the statistics of putative CA1 regions and SWRs. Only the first two sessions (sessions #1 and #2) from each subject were utilized to reduce the sampling bias.

### 2.5 Statistical evaluation

The Brunner–Munzel test and the Kruskal–Wallis test were conducted by the Python package scipy (Virtanen et al., 2020). A correlation analysis was performed by finding the rank of the observed correlation coefficient in the corresponding set-size-shuffled surrogate with an in-house Python script. The bootstrap test was also conducted by an in-house Python script.

### 2.6 Code availability

The source code is available at https://github.com/yanagisawa-lab/hippocampal-neural-trajectory-fluctuations-during-a-WM-task.

## 3 Results

### 3.1 iEEG recording and the neural trajectory in MTL regions during a WM task

We used an open dataset (Boran et al., 2020). LFP signals (Figure 1A) in the MTL regions (Table 1; *e.g.*, left hippocampal head [AHL]) were recorded during a modified Sternberg task. SWR^+^ candidates were determined from the ripple band passed LFP signals (Figure 1B) in the hippocampal regions (see Methods), and SWR^−^ candidates were defined at the identical timestamps of SWR^+^ candidates but in different trials (Figure 1). Multiunit spikes (Figure 1C) were determined using a spike sorting algorithm (Niediek et al., 2016) and provided as a part of the dataset. Based on 50-ms binned multiunit activity, GPFA (Yu et al., 2009) computed the neural trajectory (= factors) of the MTL regions by session and region (Figure 1D). Each factor was z-normalized by session and region (*e.g.*, session #2 in AHL of subject #1). The Euclidian distance from the origin O was calculated (Figure 1E).

### 3.2 Hippocampal neural trajectory correlated with a WM task

In Figure 2A, the median of neural trajectories of 50 trials was exemplified as points in the three major factor space. The optimal embedding dimension for the GPFA model was estimated as three via the elbow method (Figure 2B).

The trajectory distance from the origin (O) of the hippocampus (1.11 [1.01]; median [IQR]; *n* = 195,681 time points) was larger than those of the EC (0.94 [1.10]; median [IQR]; *n* = 133,761 time points) and amygdala (0.78 [0.88]; median [IQR]; *n* = 165,281 time points) (Figure C & D). The distance between geometric medians (*e.g.*, d(gF, gE)) in trials of the hippocampus (0.60 [0.70]; median [IQR]; *n* = 8,772 combinations) was larger than those of the EC (0.28 [0.52], median [IQR]; *n* = 5,017 combinations; *p* < 0.01; Brunner–Munzel test) and amygdala (0.24 [0.42], median [IQR]; *n* = 7,466 combinations; *p* < 0.01; Brunner–Munzel test).

### 3.3 Memory load-dependent neural trajectory distance between the encoding and retrieval states in the hippocampus

In the hippocampus, the set size (*i.e.*, the number of alphabetical letters to encode) was significantly correlated with the correct rate, response time, and trajectory distance between gE and gR (= d(gE, gR)) during the WM task.

Specifically, the correct rate of trials with set size four (0.99 ± 0.11, mean ± SD; *n* = 333 trials) was higher than that of trials with set size six (0.93 ± 0.26, mean ± SD; *n* = 278 trials; *p* < 0.001, the Brunner–Munzel test with Bonferroni correction) and trials with set size eight (0.87 ± 0.34, mean ± SD; *n* = 275 trials; *p* < 0.05; Brunner–Munzel test with Bonferroni correction) (Figure 3A; *p* < 0.001, the Kruskal–Wallis test; *p* < 0.001, correlation analysis with the correlation coefficient = − 0.20).

**Figure 3.**
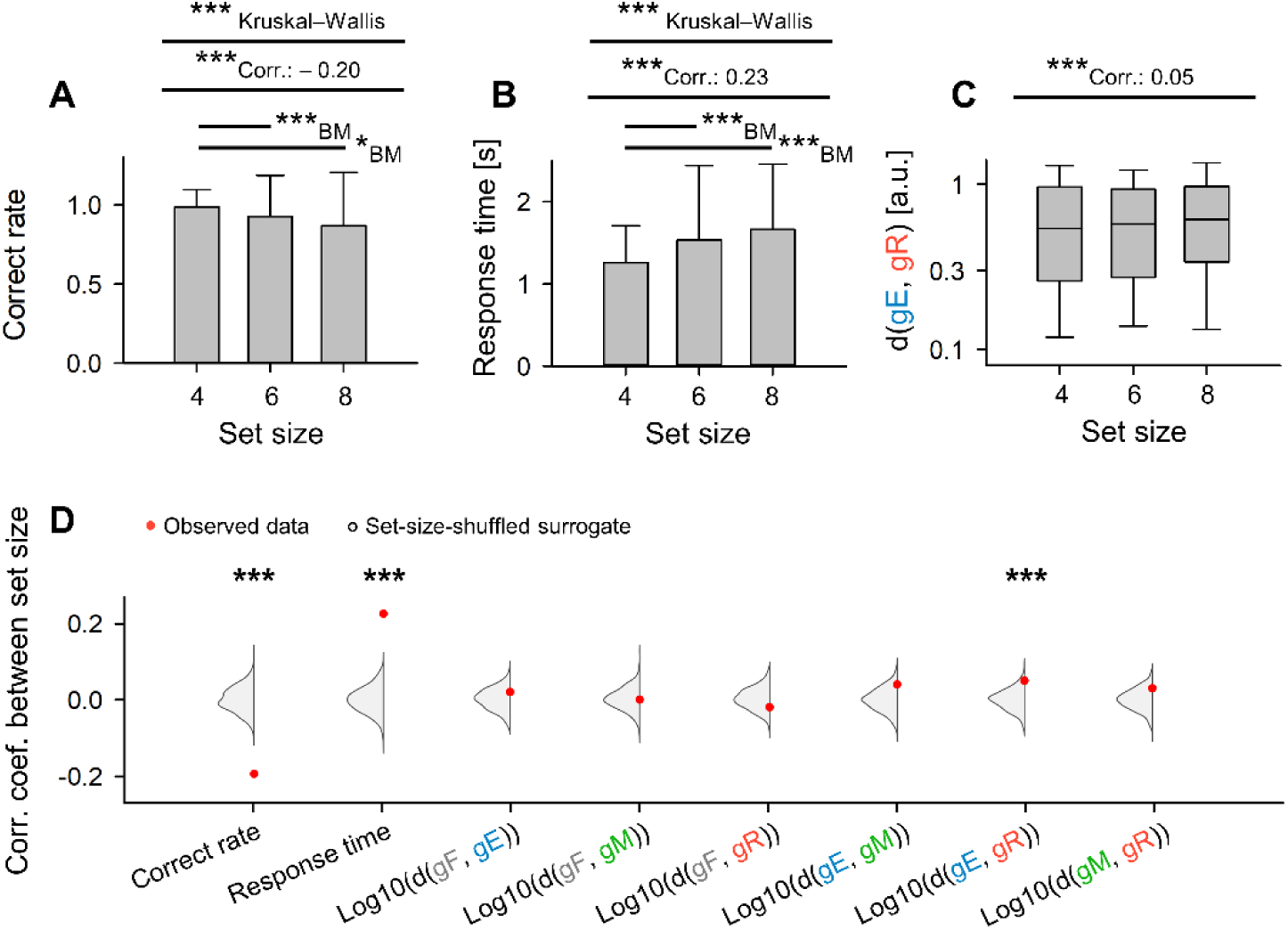
Memory load-dependent distance between the encoding and retrieval states in the hippocampus. ***A***. Set size (= the number of letters to encode) and correct rate of the WM task (\*\*\**p* < 0.001; the Kruskal–Wallis test and the Brunner–Munzel test as a post hoc test with the Bonferroni correction; correlation coefficient = − 0.20, \*\*\**p* < 0.001, correlation analysis using set-size-shuffled surrogate). ***B***. Set size and response time from probe starting time (\*\*\**p* < 0.001; the Kruskal– Wallis test and the Brunner–Munzel test as a post hoc test with the Bonferroni correction; correlation coefficient = 0.23, \*\*\**p* < 0.001, correlation analysis using set-size-shuffled surrogate). ***C***. Set size and distance between trajectory geometric medians of the encoding and retrieval phases (= d(gE, gR)) (correlation coefficient 0.05 for set size and log10(d(gE, gR)); \*\*\**p* < 0.001; correlation analysis using set-size-shuffled surrogate). ***D***. *Red* dots represent experimentally observed correlation coefficients between set size and each variable (*i.e.*, correct rate [− 0.20], response time [0.22], log10(d(gF, gE) [0.02], log10(d(gF, gM) [0.00], log10(d(gF, gR) [− 0.02], log10(d(gE, gM) [0.04], log10(d(gE, gR) [0.05], log10(d(gM, gR)) [0.03]). *Gray* violin plots show the corresponding set-size-shuffled surrogate (*n* = 1,000) (\*\*\**p* < 0.001; observed correlation coefficient vs. surrogate). Abbreviations: gF, gE, gM, gR, trajectory geometric median of trajectory during the fixation, encoding, maintenance, and retrieval phases, respectively.

Similarly, the response time of trials with set size four (1.26 ± 0.45 s, mean ± SD; *n* = 333 trials) was smaller than that of trials with set size six (1.53 ± 0.91, mean ± SD; *n* = 278 trials) and set size 8 (1.66 ± 0.80, mean ± SD; *n* = 275 trial) (Figure 3B; *p*s < 0.001, the Brunner–Munzel test with Bonferroni correction, *p* < 0.001, the Kruskal–Wallis test; *p* < 0.001, correlation analysis with the correlation coefficient = 0.22).

Furthermore, set size and the log10 of the distance between geometric medians of the encoding and retrieval phases (*e.g.*, log10(d(gE, gR))) were positively correlated (Figure 3C; correlation coefficient = 0.05; *p* < 0.001, correlation test; d(gE, gR) = 0.54 [0.70] for set size four trials, *n* = 447; d(gE, gR) = 0.58 [0.66] for set size six trials, *n* = 381; and d(gE, gR) = 0.61 [0.63] for set size eight trials, *n* = 395). However, no significant correlation was observed for the distances of other combinations of geometric medians (Figures 3D & S2).

### 3.4 Detection of hippocampal SWR from putative CA1 regions

We estimated electrodes sited at putative CA1 regions of the hippocampus where SWR events with distinct multiunit spike patterns (= spike counts per unit) were recorded compared to baseline periods. For each session and hippocampal region (*e.g.*, AHL in session #1 of subject #1), SWR^+^/SWR^−^ candidates (Figure 1) were embedded in a two-dimensional space using UMAP (Figure 4A). The silhouette score, a clustering measure, was calculated for each session and MTL region (Figure 4B & Table 2). We defined five hippocampal recording sites with an average silhouette score over 0.6 as putative CA1 regions: AHL of subject #1, AHR of subject #3, PHL of subject #4, AHL of subject #6, and AHR of subject #9 (Tables 2 & 3). Of these putative CA1 regions, the PHL of subject #4 was labeled as a seizure onset zone, while the other four sites were not (Table 1).

Next, SWR^+^/SWR^−^ candidates in putative CA1 regions were defined as SWR^+^ (*n* = 1,170) and SWR^−^ (*n* = 1,170), respectively (Table 3). The durations of both SWR^+^ and SWR^−^ were 93.0 [65.4] ms (median [IQR]; *n*s = 1,170; Figure 4C). SWR^+^ incidence increased during 0–400 ms from the probe time (Figure 4D; vs. the bootstrap sample; 95^th^ percentile = 0.42 [Hz]; *p* < 0.05). The peak ripple band amplitude of SWR^+^ (3.05 [0.85] SD of baseline, median [IQR]; *n* = 1,170) was larger than that of SWR^−^ (2.37 [0.33] SD of baseline, median [IQR]; *n* = 1,170; *p* < 0.001; Brunner–Munzel test) (Figure 4E).

### 3.5 Transient neural trajectory change in the hippocampus during SWR

The peri-SWR distance from the origin (O) was calculated in the encoding and retrieval phases (*i.e.*, eSWR^+^, eSWR^−^, rSWR^+^, and rSWR^−^) (Figure 5A). Based on the steep peak in the distance from O during SWR, we split SWR into the following three periods of events: pre SWR (= event at − 800 to – 300 ms from SWR center), mid SWR (= event at – 250 to + 250 ms from SWR center), and post SWR (= event at + 300 to + 800 ms from SWR center).

**Figure 5.**
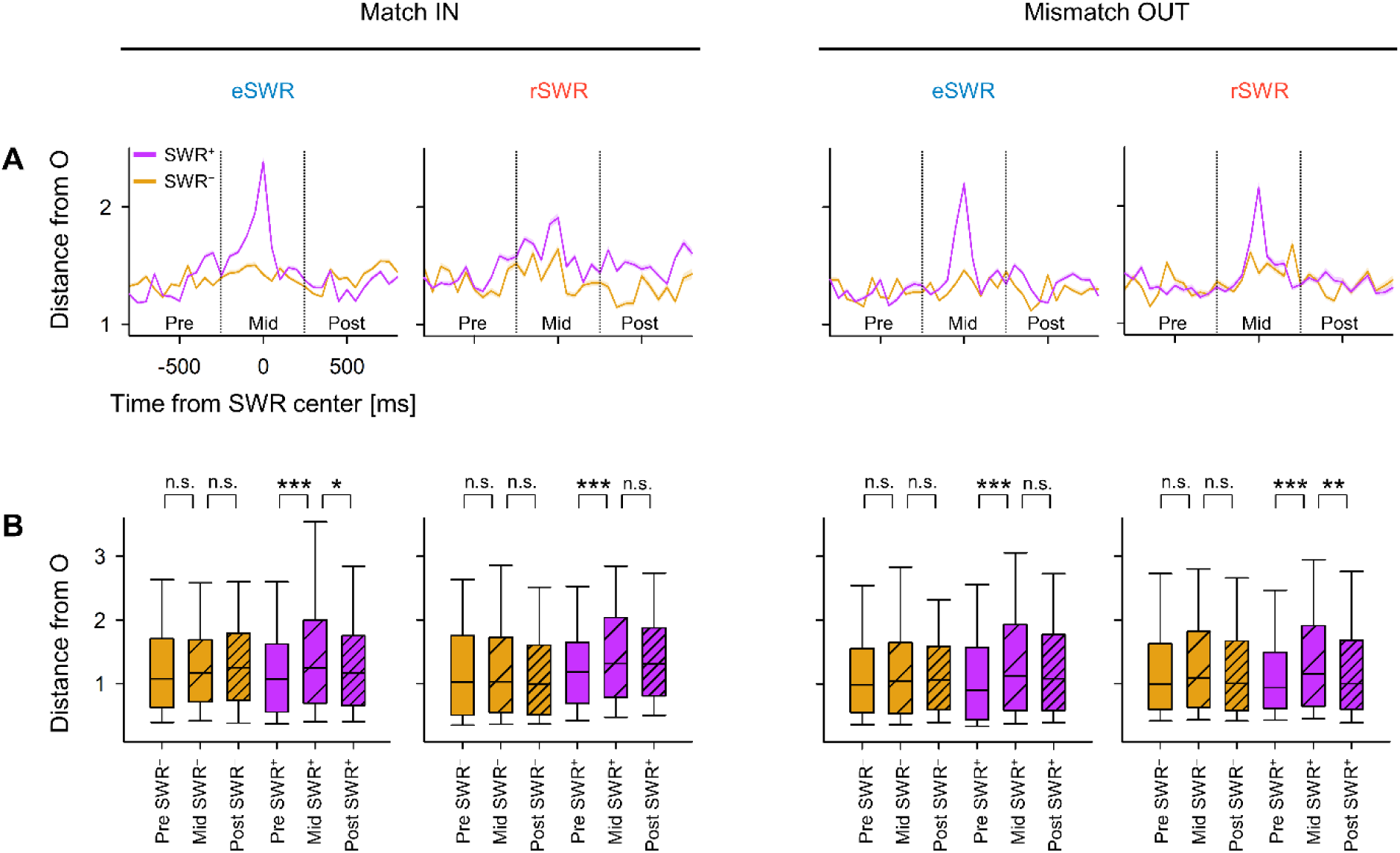
Transient neural trajectory change during SWR. ***A***. Distance from the origin (O) of peri-sharp-wave-ripple (peri-SWR) trajectory (mean ± 95% confidence interval). SWR^+^ is a putative SWR event, and SWR^−^ is a control event of SWR^+^. Based on panels in *A*, SWR events were divided into three parts: pre SWR (− 800 – − 250 ms of SWR center), mid (− 250 – + 250 ms of SWR center), or post (+ 250 – + 800 ms of SWR center). ***B***. Distance from O during mid SWR^+^ was longer compared to the corresponding pre SWR^+^ (\**p* < 0.05, *\*\*p* < 0.01, \*\*\**p* < 0.001; Brunner–Munzel test) Abbreviations: eSWR and rSWR, SWR in the encoding and retrieval phase, respectively.

The trajectory distance from O of mid eSWR^+^ (1.25 [1.30], median [IQR], *n* = 1,281, Match IN; 1.12 [1.35], median [IQR], *n* = 1,163, Mismatch OUT) was larger than that of pre eSWR^+^ (1.08 [1.07], median [IQR], *n* = 1,149, Match IN; 0.90 [1.12], median [IQR], *n* = 1,088) (Figure 5B). Similarly, the trajectory distance from O of mid rSWR^+^ (1.32 [1.24], median [IQR], *n* = 935, Match IN; 1.15 [1.26], median [IQR], *n* = 891, Mismatch OUT) was larger than that of pre rSWR^+^ (1.19 [0.96], median [IQR], *n* = 673, Match IN; 0.94 [0.88], median [IQR], *n* = 664, Mismatch OUT).

### 3.6 Visualization of hippocampal neural trajectory during SWR in two-dimensional spaces

From the previous results, it was revealed that the neural trajectory of the hippocampus “jumps” during SWR (Figure 5). Here, we visualized peri-SWR trajectories relative to the encoding and retrieval states, which exhibited memory load-dependent distance (Figure 3).

In Figure 6, the pre-, mid-, and post-SWR trajectories were plotted in two-dimensional spaces as follows. First, the peri-SWR trajectories were linearly aligned so that gE and gR were located on the x-axis. Specifically, gE was shifted toward the origin (0, 0), and gR was placed at (d(gE, gR), 0). Next, the peri-SWR trajectory was rotated around the x-axis (= gEgR axis) to be discovered on two-dimensional spaces for visualization purposes. Note that, regarding any point P (x, y), (i) the distances d(P, gE) and d(P, gR) and (ii) the angle between PgE and gEgR are preserved from those in the original three-dimensional spaces.

**Figure 6.**
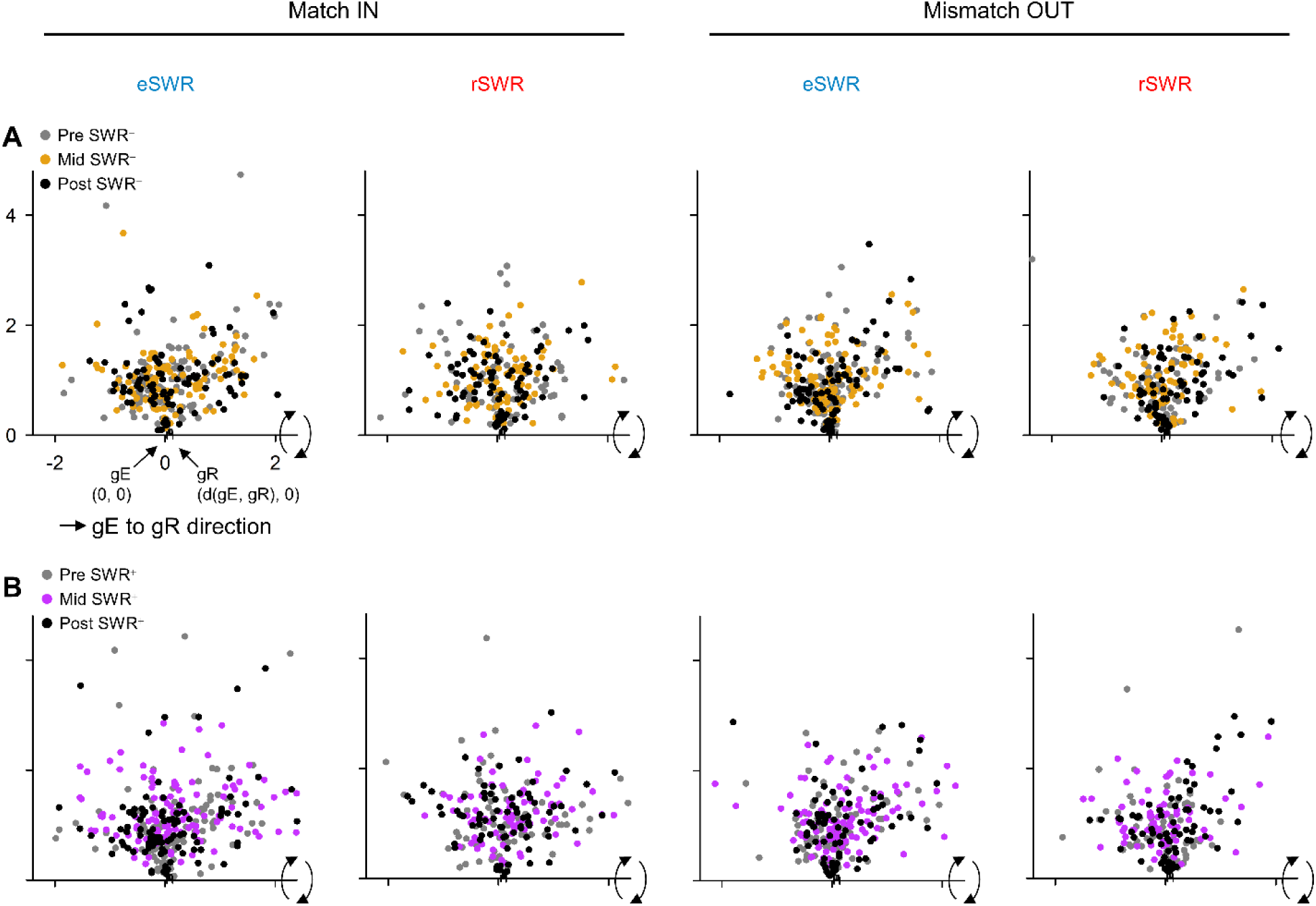
Neural trajectory coordinates during SWR aligned by the encoding and retrieval states. ***A***. ***B***. Panels show the neural trajectory coordinates during pre (− 800 – − 250 ms; *gray*), mid (− 250 – + 250 ms; *yellow* or *purple*), and post (+ 250 – + 800 ms; *black*) sharp-wave ripple (SWR^−^(***A***) or SWR^+^ (***B***)). From the left to right columns, the following events are displayed: eSWR in the Match IN task (the 1^st^ column), rSWR in the Match IN task (the 2^nd^ column), eSWR in the Mismatch OUT task (the 3^rd^ column), and rSWR in the Mismatch IN task (the 4^th^ column). The trajectory coordinates were rebased so that gE and gR were (0, 0) and (d(gE, gR), 0), respectively. Additionally, each point was rotated by the gEgR axis (= x-axis) to fit into two-dimensional spaces. d(gE, gR) were different among sessions, and their median ± IQR ranges (the medians were approximately 0.2) are shown on the x-axes. Note that regarding any point P in the two-dimensional spaces, both (i) the distance between P and gE (or gR) and (ii) the angle between gEP and gEgR (= x-axis) are preserved as those in the original three-dimensional spaces. Abbreviations: eSWR and rSWR, SWR in the encoding and retrieval phases, respectively; set size, the number of alphabetical letters for encoding.

From the scatter plot in two-dimensional spaces, we confirmed different distributions of the peri-SWR trajectory by the phases and task types, such as the longer distance between gE and mid eSWR^+^ than between gE and pre eSWR^−^, which is consistent with the result in the previous section (Figure 5B).

### 3.7 Hippocampal neural trajectory fluctuation between the encoding and retrieval states

Next, we calculated the “direction” of the trajectory relative to the encoding and retrieval states. “SWR direction” was defined based on the trajectory positions at − 250 ms and + 250 ms of SWR.

The cosine similarity between the eSWR^+^ and rSWR^+^ directions was biased toward + 1 in the Match IN task (Figure 7A) and toward – 1 in the Mismatch OUT task (Figure 7B), although the same tendencies were recognized between the eSWR^−^ and rSWR^−^ directions (Figure 7C–F).

**Figure 7.**
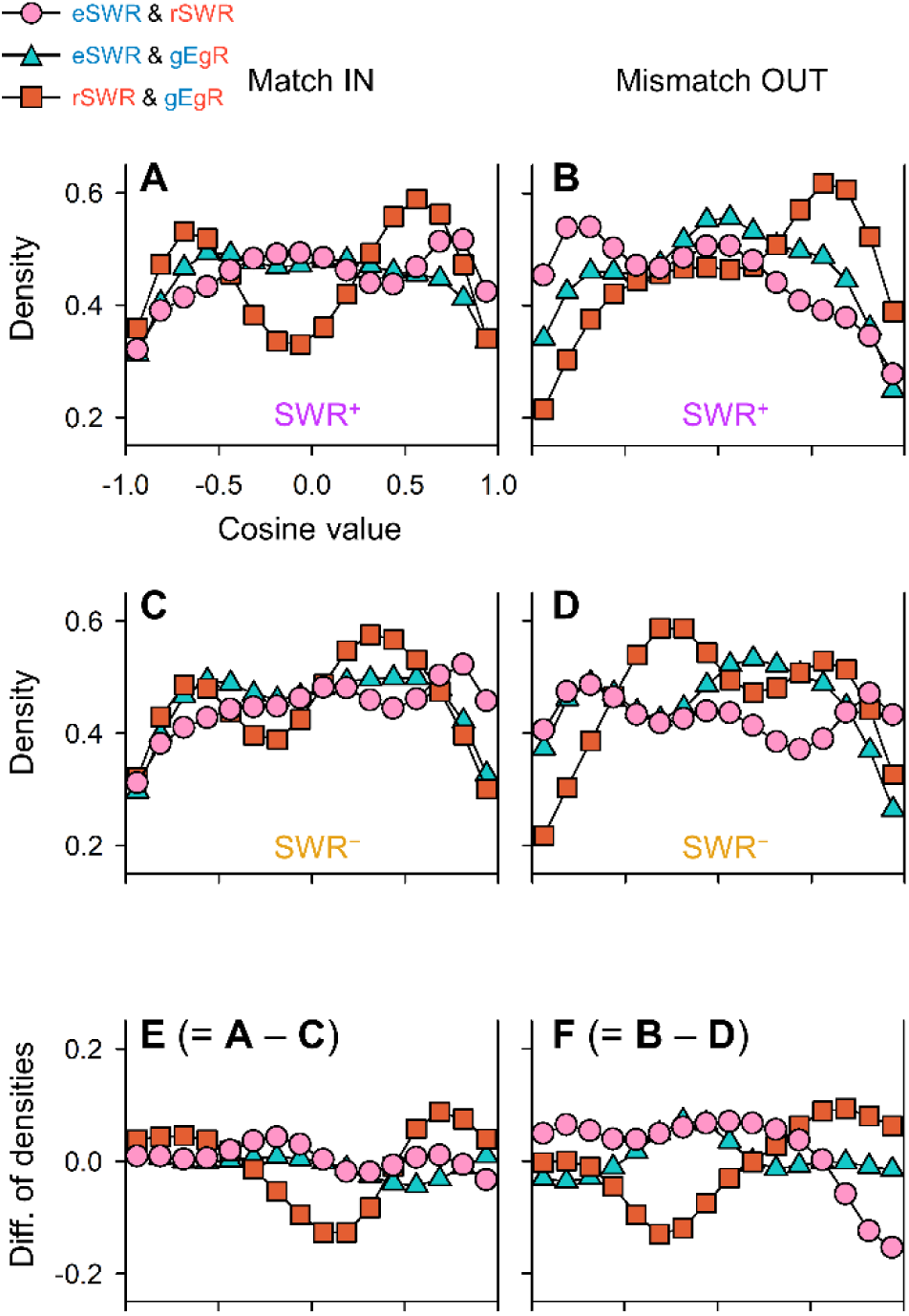
Neural trajectory directions among SWR, encoding, and retrieval states. ***A–D***. Kernel density estimation (KDE) of the distribution of cosine similarities between the trajectory directions of (i) eSWR, (ii) rSWR, and (iii) gEgR. ***E–F***. The difference between KDE values (*E* = *A* − *C*; *F* = *B* − *D*). Note the positive (/ negative) cosine similarity between eSWR and rSWR in the Match IN task (/ Mismatch OUT task) (*Pink*; *A*–*D*). Additionally, note the biphasic distribution of the cosine similarity between baseline (rSWR^−^) and gEgR (*Red*; *C*–*D*), which were shifted to the retrieval state during SWR periods (*Red*; *E*–*F*).

The cosine similarity between the rSWR^+^ direction and gEgR direction exhibited a biphasic distribution in the Match IN task (Figure 7A) but a monophasic distribution with a positive peak in the Mismatch OUT task (Figure 7B). Similarly, the cosine similarity between the rSWR^−^ direction and gEgR direction exhibited biphasic distributions, with a positive peak in the Match IN task (Figure 7C) and a negative peak in the Mismatch OUT task (Figure 7D).

Compared to rSWR^−^ directions, rSWR^+^ directions were more biased toward the retrieval state (Figure 7E & F).

## 4 Discussion

In this study, we hypothesized that hippocampal neurons express distinct representations as trajectories in low-dimensional spaces during a WM task, with a specific focus on the SWR periods. First, we embedded neural representations of the MTL regions in three-dimensional spaces (Figure 2A). The distances among trajectory geometric medians during phases were greater in the hippocampus than in the EC (entorhinal cortex) and amygdala (Figure 2E). Second, the distance between the encoding and retrieval phases (d(gE, gR)) was positively correlated with memory load (Figure 3C&D). The trajectory distance from O increased during SWR (Figure 5). Consistently, the peri-SWR trajectory exhibited a characteristic distribution in three-dimensional space (Figure 6). Finally, the hippocampal neural representation proceeded in different directions based on the encoding and retrieval phases, especially during SWR periods. In sum, these results demonstrated the involvement of hippocampal neurons and SWR in a WM task.

First, we found that the distance of the trajectory geometric medians during the four phases of the WM task was longer in the hippocampus than in the EC and amygdala, even after considering the distance from O in those regions (Figure 2C–E), indicating hippocampal participation in the WM task. These results are partly supported by previous findings of persistent firing rate increases in the maintenance phase in the hippocampus (Boran et al., 2019; Kamiński et al., 2017; Kornblith et al., 2017; Faraut et al., 2018). However, when we applied GPFA to our data, we found that the distance between the encoding and retrieval phase, d(gE, gR), was correlated with memory load (Figure 3). Overall, these results provide evidence that the hippocampus is linked to WM.

We limited our analysis to only putative CA1 regions where distinct spike unit patterns underlying SWRs were observed (Figure 4). This decision was based on the fact that SWR is time-locked to the synchronous spike bursts of interneurons and pyramidal neurons (Buzsáki, 1989; Quyen et al., 2008; Royer et al., 2012; Hajos et al., 2013), possibly ∼50 μm from the recording electrode (Schomburg et al., 2012). In fact, in our analysis, the SWR incidence increased at 0–400 ms from the probe starting time (Figure 4D), which is consistent with previous reports that hippocampal SWR occurrence increases before spontaneous verbal recall (Norman et al., 2019; Norman et al., 2021). The log-normal distribution of SWR duration and ripple band peak amplitude (Figure 4C & E) further supports the validity of the detection of physiological SWRs in this study (Liu et al., 2022). Therefore, we would have precisely captured a subclass of SWRs. One limitation is that because of the selection of channels, the increase in trajectory distance from O during SWR (Figure 5) would be biased to be greater, although it is not a critical problem in this study. In short, we were able to verify the selection of putative CA1 channels and determined SWRs, as well as the calculation of trajectory by GPFA.

Interestingly, the trajectory direction during SWR was distributed differently according to the phases (= encoding and retrieval phases) and task types (= Match IN or Mismatch OUT task) of the WM task (Figure 6). For example, the directions of eSWR^+^ and rSWR^+^ were not similar in the Mismatch OUT task but in the Match IN task. This result is reasonable because all letters should be recalled in the Mismatch OUT task, while only a part of the letters was enough to be recalled in the Match IN task. Furthermore, in the Mismatch OUT task, the probe letter was not included in the letters in the encoding phase. Thus, it might be suggested that the role of SWR in the WM task was expressed as their directions in the low-dimensional spaces.

Last, the directions of the trajectory in the retrieval phase oscillated between the encoding and retrieval states (Figure 7C & D), and the balance of such fluctuation was shifted to the retrieval state during SWR (Figure 7 E & F). These results are reasonable from the engram cell theory (Liu et al., 2012) because both the Match IN and Mismatch OUT tasks required subjects to recall the letters in the encoded phase, but the probe letter first appeared during the retrieval phase in the Mismatch OUT task. In addition, again, these results are partially consistent with previous reports that suggest the role of SWR in memory recall (Norman et al., 2019; Norman et al., 2021). Therefore, our results provide a new aspect of the hippocampal representations during retrieval periods, the neural fluctuation between the encoding and retrieval phases, and suggest the role of SWR in the retrieval phase in WM tasks.

In conclusion, this study demonstrated the involvement of the hippocampus in a WM task. We showed that hippocampal neural representations fluctuated between the encoding and retrieval states in low-dimensional spaces during a WM task. Thus, our results provide an update to the two-stage model of memory formation (Marr, 1971; Buzsáki, 1989).

## Contributors

Y.W. and T.Y. conceptualized the study; Y.W. performed the data analysis; Y.W. and T.Y. wrote the original draft; and all authors reviewed the final manuscript.

## Acknowledgments

This research was funded by a grant from the Japan Agency for Medical Research and Development (AMED) (JP19de0107001). This work was also supported in part by the Japan Science and Technology Agency (JST), Core Research for Evolutional Science and Technology (JPMJCR18A5), Exploratory Research for Advanced Technology (JPMJER1801), Moonshot R&D-MILLENNIA Program (JPMJMS2012), Grants-in-Aid for Scientific Research from the Japan Society for the Promotion of Science (JSPS) (JP20H05705, JP18H04085 and JP18H05522), AMED (JP19dm0307103, 19dm0207070h0001 and JP19dm0307008), Council for Science, Technology and Innovation (CSTI), Cross-Ministerial Strategic Innovation Promotion Program (SIP), and “Innovative AI Hospital System” (Funding Agency: National Institute of Biomedical Innovation, Health and Nutrition (NIBIOHN)).

