## Supplemental file for "Hippocampal neural fluctuation between memory encoding and retrieval states during a working memory task in humans"

### Figures

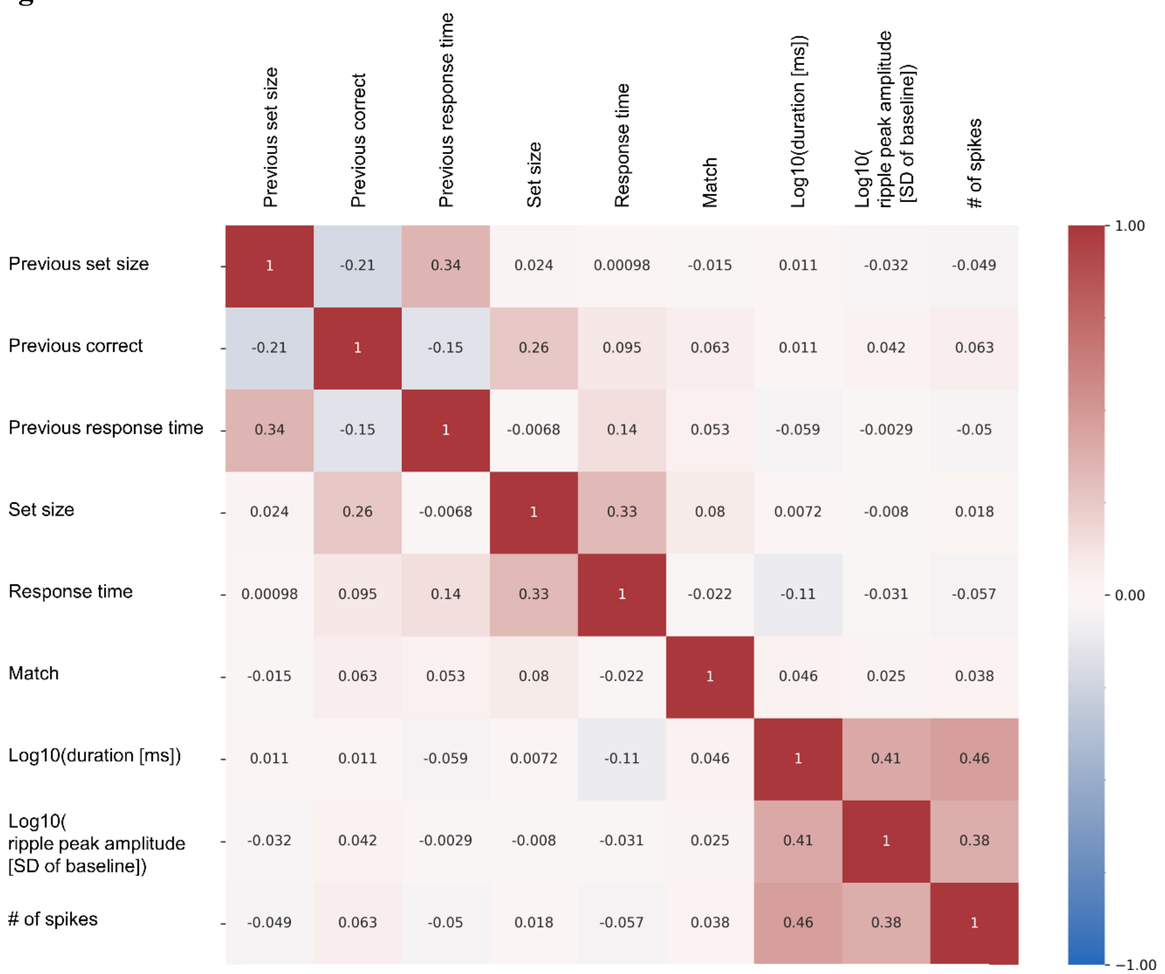


**Figure S1. Correlations among variables**

The heatmap shows the correlation coefficients among variables related to the modified Sternberg task and detected sharp-wave ripples (SWRs) parameters. Note that, the “previous correct” and set size (= the number of alphabetical letters to encode) showed the 0.26 correlation coefficient because the set size was always fixed as four after incorrect tasks. Abbreviations: previous set size, the set size of a previous trial; previous correct, incorrect (= 0) or correct (= 1) of the previous trial; match, Match IN (0) or Mismatch OUT (1) of the trial; log10(duration

[ms]), log10(duration [ms]) of sharp-wave ripple (SWR) event; log10(ripple peak amplitude [SD of baseline]), log10(ripple peak amplitude [SD of baseline]) of SWR event; # of spikes, the number of spikes during a SWR event.

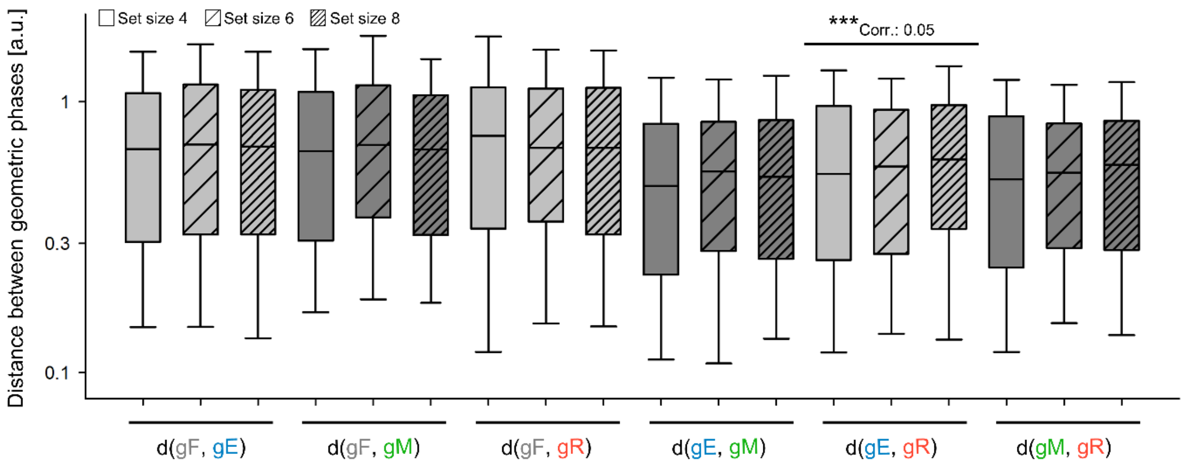


**Figure S2. Memory load-dependent distance between the encoding and retrieval states**

The figure shows distances between the four geometric phases (*i.e.*, the fixation, encoding, maintenance, and retrieval phases). Among all the combinations, d(gE, gR) showed a significant correlation with set size (= the number of letters to encode) (***$p < 0.001$; correlation analysis using set-size-shuffled surrogate).
